# Efficient cortical coding of 3D posture in freely behaving rats

**DOI:** 10.1101/307785

**Authors:** Bartul Mimicat, Benjamin A. Dunnt, Tuce Tombaz, V.P.T.N.C. Srikanth Bojja, Jonathan R. Whitlock

**Author notes:** Equal contribution (Bartul Mimicat, Benjamin A. Dunnt).

## Abstract

In order to meet physical and behavioral demands of their environments animals constantly update their body posture, but little is known about the neural signals on which this ability depends. To better understand the role of cortex in coordinating natural pose and movement, we tracked the heads and backs of freely foraging rats in 3D while recording simultaneously from posterior parietal cortex (PPC) and frontal motor cortex (M2), areas critical for spatial movement planning and navigation. Single units in both regions were tuned mainly to postural features of the head, back and neck, and much less so to their movement. Representations of the head and back were organized topographically across PPC and M2, and the tuning peaks of the cells were distributed in an efficient manner, where substantially fewer cells encoded postures that occurred more often. Postural signals in both areas were sufficiently robust to allow reconstruction of ongoing behavior with 90% accuracy. Together, these findings demonstrate that both parietal and frontal motor cortices maintain an efficient, organized representation of 3D posture during unrestrained behavior.

## Introduction

How neural circuits encode natural movement in 3D is a fundamental question in neuroscience. Though the neuronal computations involved are not fully understood, a substantial volume of work in head-fixed primates has identified key roles for posterior parietal and pre-motor cortices^1 2 3 4 5 6 7^. These efforts have produced a foundational understanding of cortical signals supporting targeted movements of individual parts of the body such as the eye or hand, with more recent studies elucidating population codes for 3D reaching trajectories and applying them to drive prosthetic limbs^8 9 10 11^. Parallel work in freely behaving rodents has demonstrated ostensibly similar functions for PPC and frontal motor areas in spatial orienting behavior^12^, movement planning^13 14 15^, and spatial navigation^16 17 18 19^. Although the rodent and primate literatures are empirically consistent, the field lacks a quantitatively precise account of how the cortex encodes naturally paced behavior in freely behaving individuals.

We therefore tracked the head and back of unrestrained rats in high-resolution 3D while recording neural ensemble activity simultaneously from PPC and the frontal motor area with which it is connected, M2^20^. By combining these methods we found the first evidence in any freely behaving model that both regions express robust population codes for 3D posture of the head, back and whole body, and that postural tuning in all animals was skewed so that fewer neurons represented poses in which the animals spent the most time. As our dataset included >1500 cells, we were able to discern that head and back representations were organized topographically over the cortical surface, and that spiking activity in PPC largely preceded M2, implying a directional flow of information across areas.

## Results

### 1D tuning curves for head, neck and back position

Eleven rats were given dual-microdrive implants in the right hemisphere, with silicon probes targeting deep (>500um) PPC and M2 (Extended Data Fig. 1, 2 and 3), areas which exhibit similar thalamic, cortical and sub-cortical connections as PPC and premotor areas across mammals^21 20 22 23 24 25^. We recorded 729 well-isolated single units in PPC and 808 units in M2 while tracking rats with a 6-camera infrared tracking system during 20-minute foraging sessions in a 2m octagonal arena (Extended Data Figs. 1, 2 and 3; Supplementary video 1). Head position for each animal was monitored with a 4-point rigid body fixed above the microdrives, while the body was tracked with three circles of retroreflective tape placed along the top of the back (Fig. 1a; Methods).

**Figure 1.**
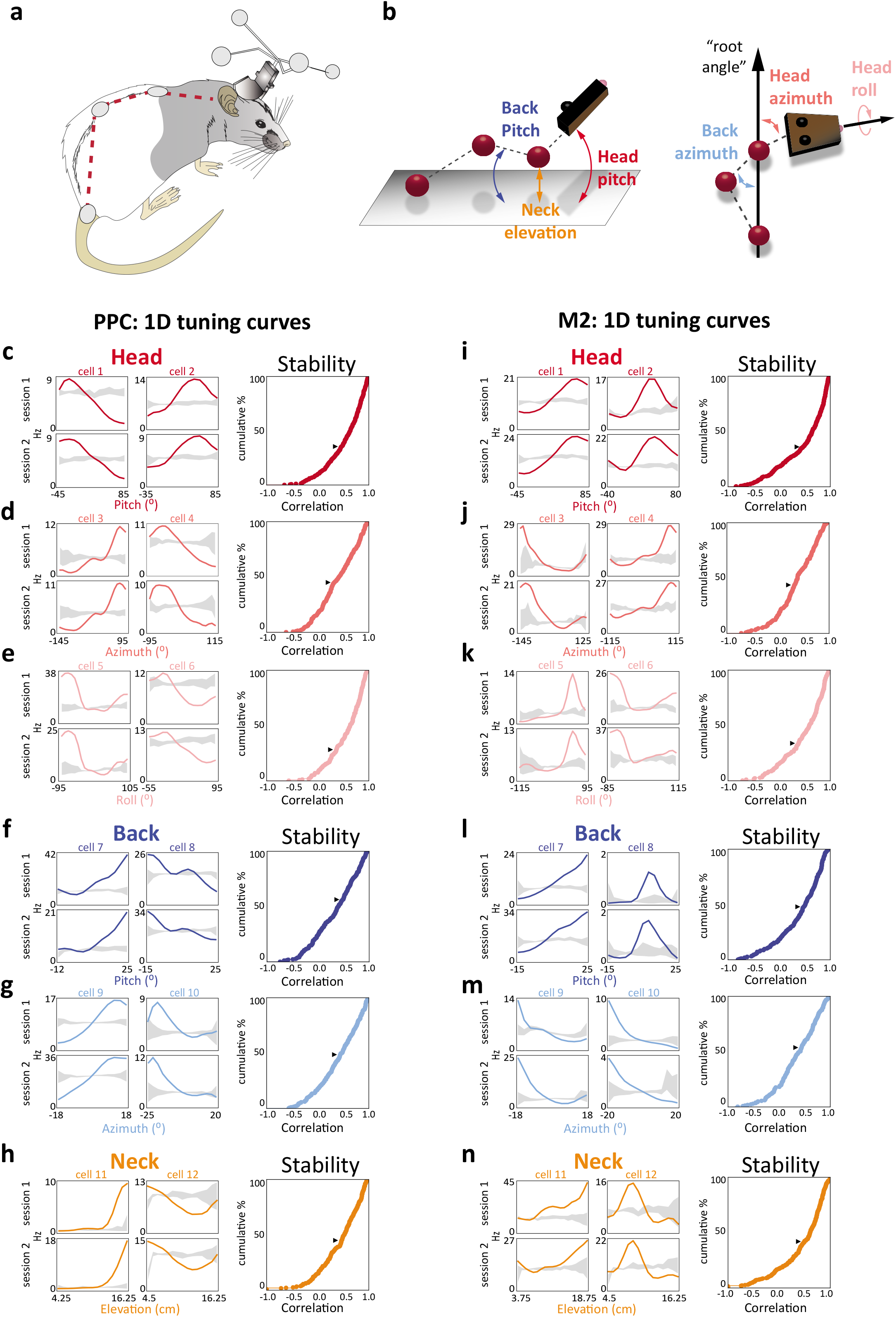
Head, back and neck position in PPC and M2: one dimensional coding of pitch, azimuth, roll and neck elevation. a, Schematic illustration of head and back tracking using retroreflective markers (Methods; Extended Data Fig. 1). b, (left) Back pitch (blue arrow), neck elevation (orange arrow) and head pitch (red arrow) were calculated relative to the floor of the arena. (right) Azimuth of the head (pink arrow) and back (light blue) were measured relative to the animal’s body axis projected onto the floor (black arrow), defined by a line anchored between the tail marker and the base of the neck. Head roll (light pink) was calculated relative to the arena floor. c-h, Examples of tuning curves for PPC cells measured in two open field sessions, with 95% confidence intervals for shuffled data shown in grey. Tuning to pitch, azimuth and roll of the head are shown (top, red), as well as back pitch and azimuth (middle, blue), and neck elevation (bottom, orange). Cumulative frequency histograms showing the stability of the tuning curves for each feature (black triangles mark the 95^th^ percentile of the null distribution; detailed results in Extended data Table 1). i-n, Same as c-h for cells recorded in M2.

This configuration allowed us to estimate Euler angles (pitch, azimuth and roll) of the head relative to the body, as well as pitch and azimuthal flexion of the back, and elevation of the neck during single unit recordings (Fig. 1b; Methods). In doing so, we found substantial numbers of cells in PPC and M2 that exhibited robust 1D tuning curves for all postural features, with peak rates often >5 standard deviations (S.D.) from the null distribution (Fig. 1c-h, i-n). The shapes of the tuning curves could extend above and below the shuffled distribution, indicating that suppression of firing was also informative of pose. The majority of cells with 1D tuning peaks exceeding the 99^th^ percentile of the shuffled distribution (Extended Data Fig. 4a,b) were stable across recording sessions (mean of 56.4% in PPC, 57.8% in M2, Fig. 1c-h, i-n; Extended data Table 1), and postural tuning in lighted sessions was stable in darkness (Extended Data Fig. 5).

**Figure 4.**
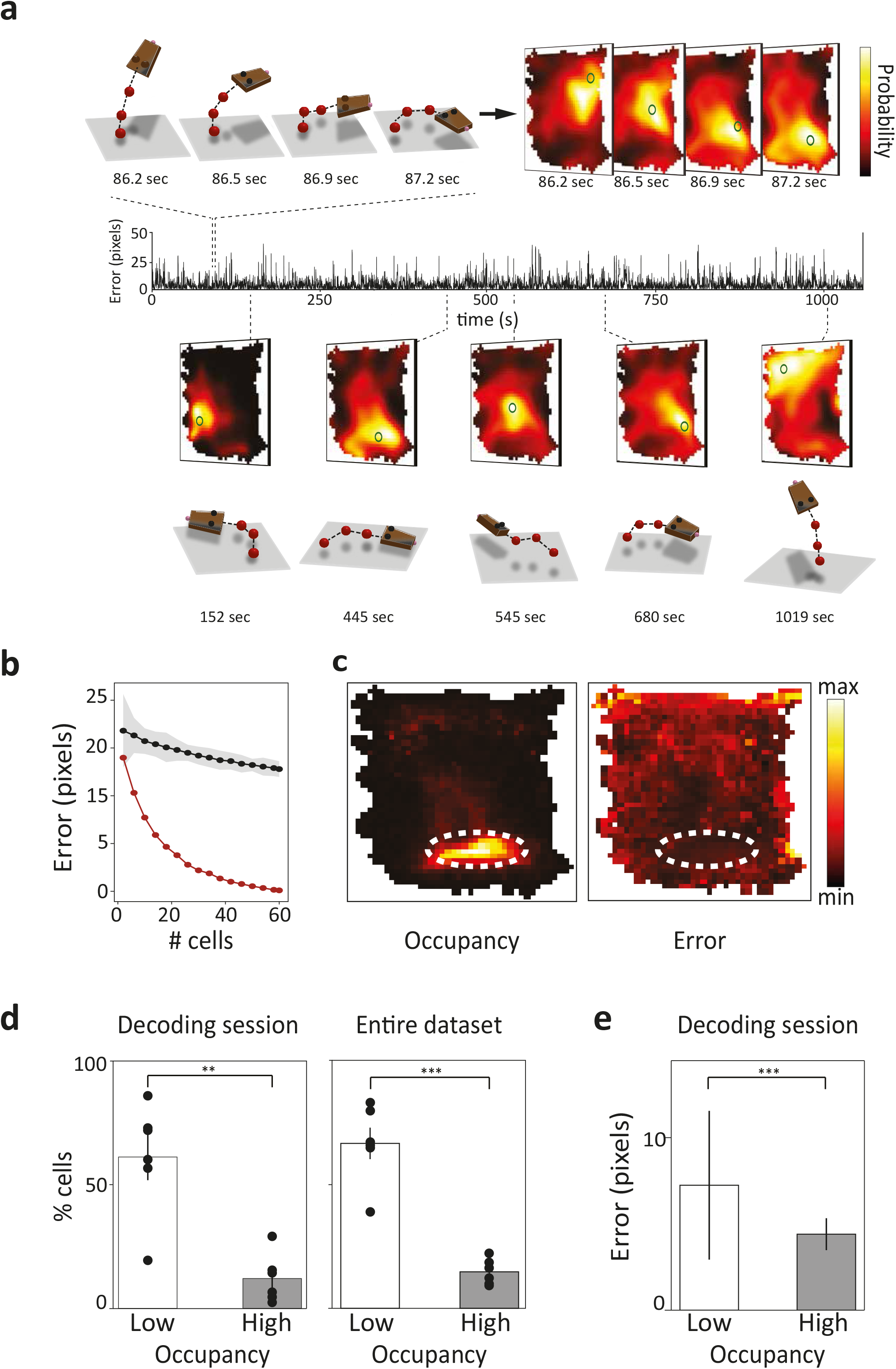
Ensemble decoding of pose from PPC and M2 reveals a non-uniform distribution of tuning in the population. a, (top, left) Four snapshots taken within 1s as the animal came down from rearing and bent rightward. (top, right) Frames from corresponding decoded maps illustrate the log posterior distribution of the animal’s posture estimated from ensemble activity in PPC and M2. Actual pose is marked with a green “O”, while the prediction is color coded yellow-to-black. (middle) Timeline indicating error, in Isomap pixels, taken from a 20 minute recording session. (bottom) Five examples of distinct poses throughout the session (time points marked underneath) and their attendant Isomaps showing real and decoded pose. b, Decoder accuracy as a function of the number of cells (red dots), with the null distribution above (black dots). Shaded area indicates ±3 S.D. c,(left) Cumulative occupancy on the Isomap shows the animal spent the majority of the session in the low centre of the map, corresponding to postures during foraging on all fours. The dashed yellow oval contains the “high occupancy” area in which the animal spent >50% of the session. d, (left) Decoder error of the posture map was significantly higher in low occupancy areas compared to high. (right) The percentage of cells encoding each postural feature (black dots) was significantly higher in low *versus* high occupancy areas. Bar graphs indicate the mean ± SEM. e, Same analysis as the right panel of d, but for all 11 animals in the study.

### PPC and M2 encode 3D pose

We found that many cells in PPC and M2 responded to conjunctions of Euler angles for the head and back, and in some cases were specific to the posture of the whole-body (Fig. 2a and c; Supplementary Videos 2-6). Since many aspects of posture and movement co-vary naturally, we disambiguated features best explaining spiking activity using an unbiased generalized linear model (GLM, Methods). The model considered egocentric pose variables and their derivatives, as well as allocentric features including head direction, running direction and spatial location. Covariates that best explained neural firing were selected using a forward-search approach, in which features were added until model performance no longer improved 10-fold cross validation^26^ (Methods).

**Figure 2.**
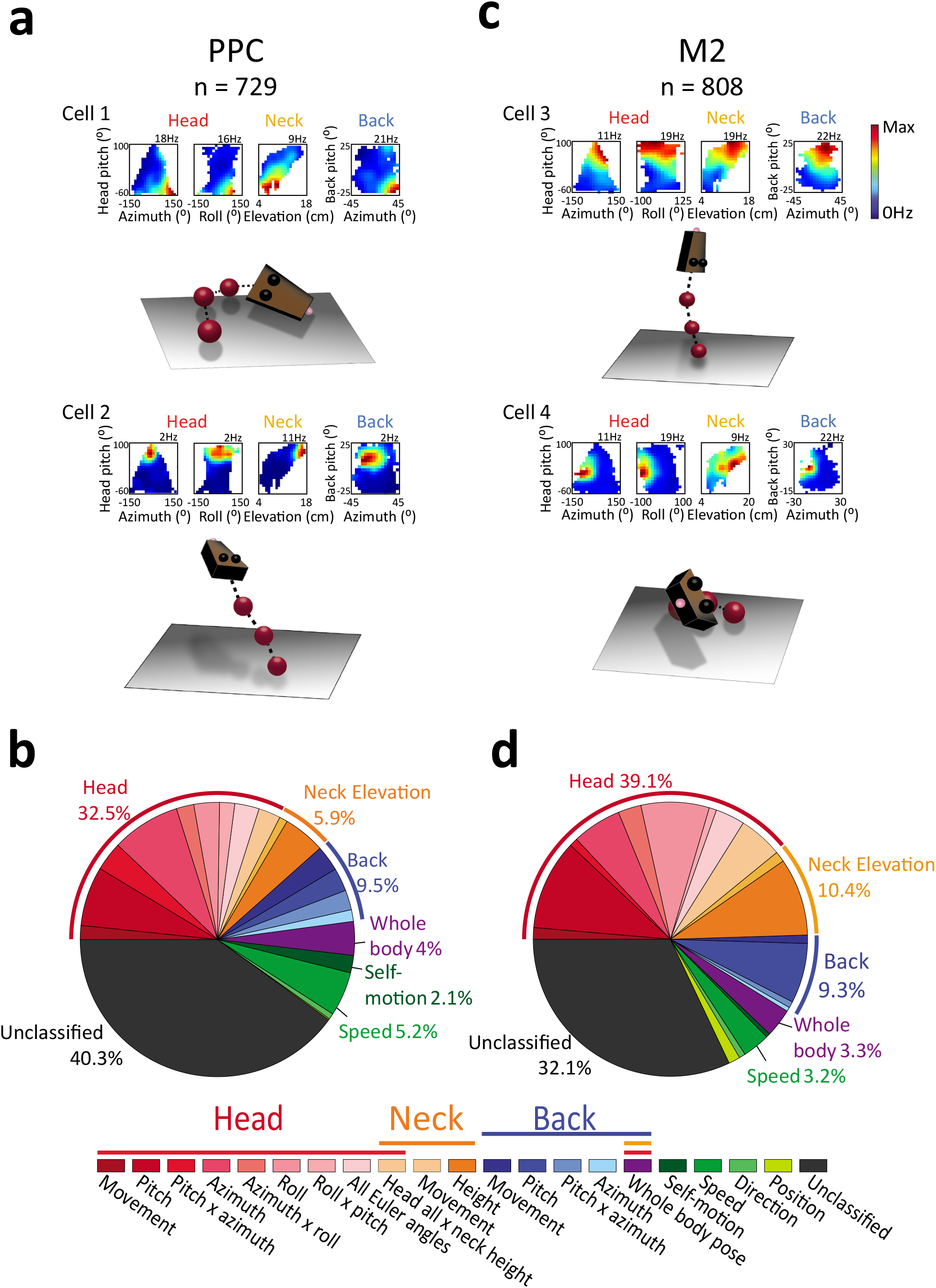
Parietal and motor cortices are tuned to combinations of head, back and neck position. a, Two example PPC cells showing tuning to combinations of head, back and neck positions; the conjunctive representations appear as single firing fields in 2D rate maps (top). Each rate map corresponds to a combination of Euler angles of the head, back, and neck elevation. (below) 3D rat models depict the postures to which each cell was tuned. Cell 1 was selective for whole-body flexion to the right plus rightward head roll; Cell 2 fired during rearing, driven primarily by the interaction of head pitch x neck elevation. b, The distribution of behavioral tuning in 729 PPC cells as determined using a generalized linear model. The largest fractions of cells in PPC were driven mainly by head position, though significant fractions of cells were tuned to neck height, back position, and whole-body postures (see color-coded legend below and Supplementary Table 2). Running speed and self-motion were also represented in the population, though much less than pose. c, Examples of postural tuning in M2 cells. Cell 3 (top right) was especially sensitive to vertical behaviors, with the head, back and neck raised; cell 4 was tuned to leftward head roll and back flexion during sharp turns. d, Distribution of coding properties across 808 cells in M2. As in PPC, head position alone accounted for nearly 40% of the cells in M2, followed by neck elevation (10.4%) and back position (9.3%).

The GLM identified 437 of 729 PPC cells showing significant tuning to at least one egocentric or allocentric variable (Extended data Table 2). The largest fraction of cells (n = 237, 32.5%) were driven by postural features of the head, such as pitch, azimuth or roll individually, interactions between Euler angles (e.g. pitch x azimuth), conjunctions of 3D head pose and neck height, and head movement (Fig. 2b). The next largest groups of cells were tuned to back posture or movement (n = 69), and neck elevation (n = 43; Fig. 2b, Extended data Table 2). A smaller number of cells exhibited “whole body” postural tuning, driven conjointly by head, neck and back position (n = 29, 4.0%, *Z* = 7.9, P < 0.001; large-sample binomial test with expected *P*_0_ of 0.01).

In contrast to previous studies on rat PPC^14 15^, we found modest fractions of cells showing tuning to running speed (n = 38, 5.2%, *Z* = 11.2, P < 0.001) and self-motion (n = 15, 2.1%, *Z* = 2.7, P < 0.01; Fig. 2b; Extended data Table 2), raising the possibility that those studies, which used 2D tracking, failed to resolve movement types from the 3D postures that attend them (Extended Data Fig. 6). Allocentric features such as head direction, running direction (combined n = 5, 0.7%; *Z* = −0.67, P > 0.85) and spatial position (n = 1, 0.1%, *Z* = −2.2, P > 0.99) were not significant in the data set.

The 808 cells recorded in M2 showed tuning to similar variables as PPC, though in different proportions (Fig. 2c). For example, half (n = 400, 49.5%) the cells encoded 3D head posture, head movement or neck elevation (Fig. 2d, Extended data Table 2). The fractions of M2 cells tuned to posture and movement of the back (n = 75, 9.3%), as well as whole-body pose (n = 27, 3.3%) were on par with those seen in PPC. As in PPC, a small but significant fraction of cells was tuned to running speed (n = 26, 3.2%, *Z* = 6.2, P > 0.001), while selfmotion tuning (n = 4, 0.5%; *Z* = −1.3, P > 0.95; Extended data Fig. 7), direction (n = 6, 0.7%; *Z* = −0.6, P > 0.8), and location (n = 11, 1.4%; *Z* = 0.86, P > 0.15) were not significant in M2.

We were struck that our statistical model indicated that a large majority of cells in PPC and M2, a motor area, were driven by posture (46.2% in PPC; 58.7% in M2) as opposed to bodily movement (5.6% in PPC; 3.6% in M2), and sought to confirm the result. Since posture and movement often co-varied, we split recording sessions for individual cells from PPC and M2 on the basis of high and low movement velocity for specific features (e.g. head pitch), and examined how this affected the tuning curves. We found that tuning curves for pose remained virtually identical regardless of movement, while tuning curves for movement varied unreliably when split by posture (Extended data Figure 8), confirming that postural tuning was indeed expressed independently of movement.

### Spatial and temporal organization of tuning in PPC and M2

Microstimulation and electrophysiological recordings in several mammalian species have identified topographical organizations in parietal and motor areas controlling movements for many parts of the body, including, the face, mouth, head, arm and trunk^27 28 29 30 31 32^. We therefore assessed whether postural features identified by the GLM were organized anatomically, and found significant gradients for the head and back across the surface of M2 and PPC. Specifically, head representation in M2 was concentrated at anterior (χ^2^(4) = 57.1,P < 0.001; Yates corrected χ^2^ test; Fig. 3a) and medial (χ^2^(4) = 110.6, P < 0.001) locations, while back posture was represented more densely at the posterior (χ^2^(4) = 98.1, P < 0.001) and lateral (χ^2^(4) = 105, P < 0.001) poles (Fig. 3a, b). Back-dominated sites in M2 were contiguous with the most anterior and medial sites in PPC, which themselves showed the strongest coding for the back (χ^2^(3) = 29.9, P < 0.001, anterior-posterior gradient; χ^2^(4) = 12.5, P < 0.05, medial-lateral). Head posture was represented predominantly in the posterior-lateral regions of PPC (χ^2^(4) = 47.5, P < 0.001, anterior-posterior; χ^2^(4) = 52.4, P < 0.001, medial-lateral), producing a coarse mirroring of head and back representations across PPC and M2 (Fig. 3a).

**Figure 3.**
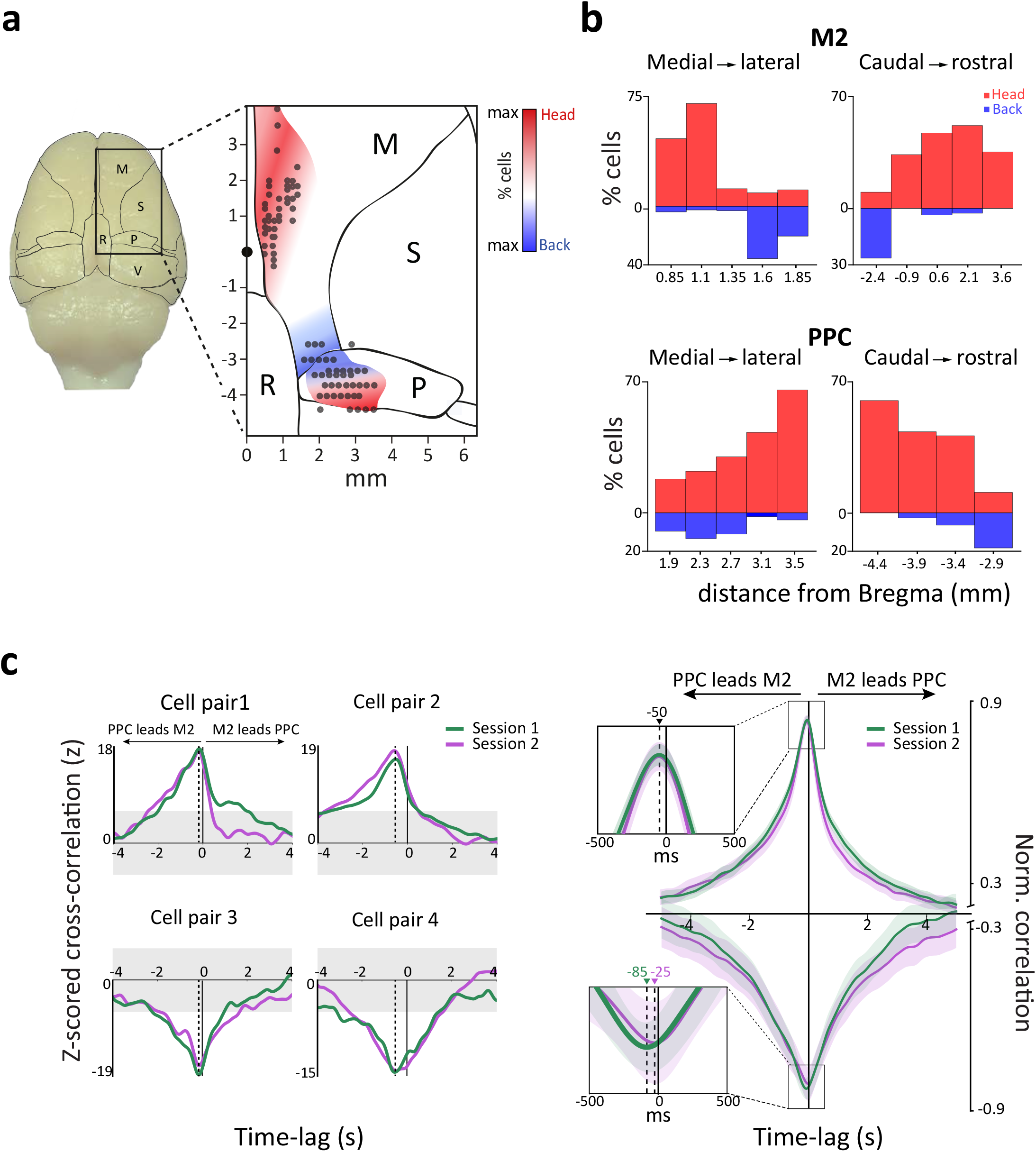
Head and back posture are organized topographically across areas, and spiking activity in PPC leads M2. a, Dorsal view of the rat brain with anatomical boundaries illustrating primary and secondary motor cortices (M), somatosensory cortices (S), retrosplenial cortex (R), posterior parietal cortex (P), and primary and secondary visual cortices (V). The magnified view (right) shows recording locations (grey dots), and colored shading indicates the functional-anatomical distribution of tuning to the head (red) or back (blue). b, Percentages of cells in M2 (top) and PPC (bottom) driven by head and back position. In all cases the actual distribution tuning differed significantly from theoretical distributions that assumed a constant proportion of tuned cells across bins. c, (left) Four examples of cell pairs in PPC and M2, each showing stable z-scored cross-correlations over two recording sessions with spiking activity in PPC preceding M2. Shaded areas indicate ±6 S.D. of the shuffled data. (right) The normalized cross-correlation for all cell pairs shows a negative peak for PPC relative to M2 for both positive and negative correlations across behavioral sessions. Shading indicates the 99% confidence interval of the mean.

Previous recordings in primates suggest that PPC and frontal motor areas comprise an anatomically extended network^33^ for planning and decision making^34 35^, but few studies have succeeded in recording spikes from both areas simultaneously. We therefore asked whether a temporal correlation structure existed between spikes recorded simultaneously in PPC and M2 (n = 5 rats). Having screened for inter-regional cell pairs whose signal crosscorrelations within a ±5 second window exceeded chance (Fig. 3c; Methods), we found 1017positively and 182 negatively correlated cell pairs in one recording session, and 758 positively and 141 negatively correlated cell pairs in a second session from the same day. In both sessions, the majority of correlated pairs (60-62%) were led by PPC, with the normalized correlation of positive pairs peaking at −50ms relative to M2 (bootstrapped 99% C.I. for session 1: −113 to +8ms; session 2: −112 to +7ms), and negative correlations peaking at −85ms (bootstrapped 99% C.I.: −191 to +16ms) and −25ms (bootstrapped 99% C.I.: −129 to +74ms) for sessions 1 and 2, respectively (Fig. 3c). The number of cell pairs fell too low for comparison when split by tuning properties, but the data on the whole are consistent with an asymmetric information flow in which PPC leads M2.

### Ensemble decoding and the non-uniformity of postural tuning

Although we observed robust postural correlates for single cells in PPC and M2, this did not answer whether population activity was sufficiently informative to reconstruct ongoing behavior. To address this we first reduced the behavioral dataset from six dimensions (3 axes for the head, 2 for the back, and one for neck elevation) to two using Isomap^36^, a dimensionality reduction algorithm that approximately preserves local distances between linked features. With this approach, we mapped posture for the head, back and neck onto a single 2D surface, or “posture map”, with each pixel corresponding to a particular bodily configuration (Extended data Fig. 9). Neighboring pixels on the map correspond to similar postures, while dissimilar postures are farther apart. Using a session with 37 PPC and 22 M2 neurons recorded simultaneously, we trained a uniform prior decoder using 10-fold crossvalidation to predict the animal’s position on the posture map, with maximal likelihood depicted by the color of the pixels (Fig. 4a; Supplementary videos 7-8). Decoder accuracy was highest with the full complement of cells (Fig. 4b), declined to 50% with 20 cells, and reached chance levels with 6 cells. Extrapolation analysis estimated that 177 cells with similar selectivity would be required to reach maximum decoding accuracy (not shown).

We noted that the cumulative occupancy of the 2D posture map was dominated by the area corresponding to the animal standing on all 4’s with the head lowered (i.e. foraging; Fig. 4c, left). Further analysis showed that a significantly larger fraction cells were tuned to postures in which the animal spent less time, while fewer cells encoded the postures the animal expressed most (t(10) = 4.82, P<0.01, Welch’s t-test; Fig. 4d, left). We repeated the analysis for our entire dataset and found the same pattern of tuning for all postural features across animals (t(10) = 7.74, P<0.001; Fig. 4d, right, Extended data Fig. 10), suggesting that the cells’ receptive fields were distributed optimally according to occupancy. Despite the low density of neural representation in the area of high occupancy, decoder error remained low. In fact, the decoder performed significantly better (t(74) = 6.21, P<0.001; Fig. 4e) in the region with >50% occupancy, despite being represented by ~15% of the ensemble. To our knowledge, such a non-uniformity of representation has not been described previously for positional coding of other effectors, or in other sensorimotor systems.

## Discussion

We recorded single units from parietal and frontal motor cortices while tracking freely moving animals, revealing that cell populations in both regions robustly encode 3D posture, and in far larger proportions than other features of behavior. By measuring neural signals encoding the dynamic position of the head and trunk during whole-body movement, these data complement and extend years of detailed study on the positional coding of individual effectors in stationary animal^4 5 37 38^. The data directly support the notion that PPC is a key cortical substrate for the continual awareness of the spatial configuration of the body, or “body schema”, postulated over 100 years ago ^39^, and suggest that frontal motor areas may also play a role.

The fact that tuning for posture vastly outstripped that for movement in PPC and M2, a motor area, was surprising and can be interpreted in different ways. One reason why postural tuning may have figured so prominently in our dataset was that the animals were freely behaving, in which case the brain must solve a host of kinematic problems not encountered when the axes of movement are limited by behavioral constraint. The imbalance of postural and movement signals in cortex also raises the possibility that a functional division of labor exists, at least partially, between cortical and sub-cortical motor systems. On this view, parietal and higher motor areas encode the positioning of the body and goals^40 41 42^, while descending pyramidal tract pathways, the basal ganglia, and brainstem motor nuclei control muscle synergies directly, and would therefore provide more robust readouts of the kinetics, force and vigor of movement. Piecing together the timing and relative contributions of each of these systems to unrestrained 3D behavior is a new and exciting prospect that will require additional study. Further work will also be required to determine whether postural tuning is expressed throughout cortex during unrestrained behavior, akin to locomotor state or speed^43 44 45^, or occurs mainly in areas contributing to the production of movement.

Our findings also have implications for previous studies of speed and self-motion tuning in PPC in freely behaving rodents, which may likely have overestimated these features due limitations inherent to 2D tracking of head-mounted LEDs^14 15^. This approach renders the animal as a point moving on a flat surface, which conveys essentially no information to distinguish vertical movements, head roll or contortions of the body. Though effective in revealing how the brain maps the physical world^46 47^, this approach is ill-suited to resolve individual behaviors animals employ, for example, when acquiring environmental inputs needed to generate such maps in the first place^48 49 50 51 52 53^. Our statistical model indicated that self-motion tuning was nevertheless found in a small but significant fraction of cells specifically in PPC, still supporting a role for encoding bodily movement through extended space. The high sensitivity of PPC and M2 cells to the angle of the head relative to the body could also potentially influence population responses in head-fixed preparations, including virtual reality paradigms^54^, though we acknowledge that several task-related computations are not likely to be explained by ongoing posture ^55 56 57^. Regardless of the nature of the task, our findings highlight the need for close behavioral monitoring in tasks seeking to distinguish cognitive functions from physical behavior.

We also found an underlying spatial distribution of postural tuning for the head and back, and a temporal organization of spiking activity across PPC and M2. The roughly symmetric organization of head and back representations that we describe matches the general organization of sensory and motor areas described in classic cortical mapping experiments^31^. To our awareness, ours is the first study to chart a functional organization of behavioral tuning within the rodent PPC and, further, across PPC and M2. Though we did not test explicitly whether postural representations were organized temporally across regions, our cross correlation analysis supports the notion that PPC and M2 in rats, like monkeys, comprises an anatomically extended network ^33 34^ that is organized hierarchically^58^, or at least temporally, with PPC operating upstream from M2.

A striking finding complemented the ubiquity of postural coding in our dataset: the arrangement of tuning peaks was neither uniform across the distribution of postural occupancy, nor did it mirror it (Fig. 4c-e, Extended data Fig. 4). On the contrary, tuning curves were likelier to have significant peaks away from the most occupied bins, a feature that permitted accurate decoding of postures in which the animal spent less time. Likewise, the small faction of cells encoding the longest-expressed poses was also sufficient for low-error decoding. This cost-reducing property, observed here in associative cortices, raises the question of optimal coding strategies and their potential metabolic benefits in other systems. Previous theoretical works have considered optimal coding strategies in sensory systems^59 60^, postulating that the range of the stimulus spectrum visited most frequently should be encoded by more cells with narrower tuning widths, but this did not appear to be the case in our data. Ganguli & Simoncelli (2014) suggest that lowering the gains of neurons tuned to the low end of the stimulus range, or increasing their number at the high end, could be metabolically efficient, though this may incur a cost on the quality of coding. The tuning scheme we observed maintained a high fidelity of coding for both long- and shortlived postures, and would bring metabolic savings, making it both precise and efficient. An accurate representation of a default-state postural stance should demand little expenditure of neural and energetic resources.

It is very intuitive that posture would have a prominent representation in cortex, whether it is for movement planning, or as a substrate for learning postural sequences to perfect an action, such as a tennis serve. The dense representation of head position relative to the body would also serve to bring environmental inputs detected by the sense organs into immediate register with the body, facilitating rapid orientation and appropriate behavioral responses. The extent to which postural signals described here are integrated with these features, particularly in frontal cortex ^32^, remains to be determined.

## Methods

### Subjects and electrode implantation

Experiments were performed in accordance with the Norwegian Animal Welfare Act and the European Convention for the Protection of Vertebrate Animals used for Experimental and Other Scientific Purposes. Neuronal activity was recorded from 9 male and 2 female Long-Evans rats (3-5 months old, 400-600 g at implantation and testing) with chronically implanted dual microdrives. Mounted on each microdrive was a silicon probe (NeuroNexus Inc., MI, USA; custom design based on A8x1-tet-2mm-200-121). One probe targeted PPC (-3.8 to −4.25mm AP, centre shank 2.7mm ML) and one targeted M2 (centre shank +0.5mm AP, 0.7mm ML). Each probe had eight 55μm wide, 15μm thick, 5mm long shanks, with four 160μm^2^ iridium recording sites configured as a tetrode at each tip; inter-shank spacing was 200μm. At surgery, animals were anesthetized in a ventilated plexiglass box with 5% isoflurane vapour, and maintained on 1.0-2.5% isofluorane for the duration of surgery. Body temperature was maintained at 37 °C with a heating pad. Once unconscious, animals received s.c. injections of analgesic (Metacam 2.5mg/kg weight, Temgesic (buprenorphine) 0.05mg/kg). Local anaesthetic (Marcain 0.5%) was injected under the scalp before making the incision. The skull was then exposed, rinsed and sterilized using 0.9% saline and 3% hydrogen peroxide. A high-speed dental drill with 0.8mm burr was used to drill holes for skull screws and craniotomies over PPC and M2. Bone-tapping stainless steel screws were inserted securely into the skull, with a single screw serving as the ground wire and reference for both drives. The microdrives were housed in a single 3D printed unit that was lowered with probes targeting the right hemisphere. The drive unit was cemented to the skull, and probes were lowered into the brain, and craniotomies were filled with a 70-30% mixture of mineral oil and low-melt bone wax (Sigma-Aldrich Norway AS, Oslo, Norway). After surgery rats awoke in a 32 °C heated chamber, were returned to their home cage, and later administered post-operative analgesics. Rats were housed individually in Plexiglas cages (45 × 44 × 30 cm) in a humidity and temperature-controlled environment, and kept on a 12 hr light/12 hr dark schedule. All training and testing occurred in the dark phase. Experiments were performed in accordance with the Norwegian Animal Welfare Act and the European Convention for the Protection of Vertebrate Animals used for Experimental and Other Scientific Purposes.

### In vivo electrophysiology and behavior

Single units, LFP or raw signal from PPC and M2 were sampled with a Digital Lynx 4SX recording station (Neuralynx Inc. Montana, USA). Rats were connected via AC-coupled unity-gain operational amplifiers above the head (via H32-to-HS36 custom head stages, Neuralynx Inc.), which connected to a motorized commutator above the arena. Elastic string was used to counterbalance weight and excess cable length, allowing animals to move freely in the recording arena.

Silicon probes were lowered in 50μm steps while the rat rested on a towel in a flower pot on a pedestal. Turning stopped when well-separated units appeared, typically between 500 and 1800μm. Data collection started when signal amplitudes exceeded ~4 times the noise level (r.m.s. 20-30μV) and units were stable for > 3hr. After recording at a given depth, PPC and M2 probes were lowered to obtain new cells; probe depths typically differed by 100μm or more between recording days.

Behavioral recordings were performed as rats foraged for crumbs of chocolate, cereal or vanilla cookies thrown randomly into an octagonal, black open-field arena (2 × 2 × 0.8 m), with the animals oriented by extra-arena room cues. Recordings began once the animals achieved routine, complete coverage of the arena. Recording sessions typically lasted just over 20 minutes (average length 1335s, total range of 610-2467s). Rats rested a minimum of 1 hour in their home cage between runs.

### Spike sorting and analysis of firing rates during behavior

For the first 6 rats, spike data were clustered automatically with KlustaKwik (http://klustakwik.sourceforge.net; Harris et al., 2000), then sorted manually using graphical cluster-cutting software as described in Fyhn et al., 2004. For the remaining 5 animals, raw signal Neuralynx files collected on the same recording day were converted to binary format and concatenated, such that the identity of isolated units could be preserved across sessions. Spike sorting was performed offline with Kilosort^61^, followed by manual curation in Phy (C. Rossant, https://github.com/kwikteam/phy). Clusters were merged or separated based on waveform similarity, spike rate auto- and cross-correlation and masked cluster quality measures. Clusters with ≥ 2% violation of a 2ms inter-spike interval were discarded. After spikes were split into individual sessions for each cluster, only cells with >100 spikes per session were kept for further analyses.

### 3D tracking and model assignment

For tracking the head, four 9mm retroreflective markers (ca. 6cm apart) were affixed to a rigid body clamped above the recording drives. For the back, three 9mm circular cut outs of retroreflective tape (3M) were each affixed to shaved locations at the shoulder blades, the hunch of the back, and above the root of the tail. Position data were recorded at 120 fps using a 6-camera infrared recording system (OptiTrack, Oregon, USA), and registered using optical motion capture software (Motive, version 1.8, 1.9 or 1.10.2; OptiTrack).

For 7 of 11 animals (tracked with Motive versions 1.9 or 1.10.2), labeling of individual markers was performed with the in-built labeling functions in Motive on raw.tak files. Specifically, the 3D data was first deleted. Then, a rigid body was constructed from the four head markers (in the same way across animals), and the three body markers were kept in a separate marker set. Both marker assets were then dragged onto the take and the “reconstruct and autolabel” function was applied. Depending on the results, errors in marker assignment were corrected and the unlabeled points were hand labeled. For each individual marker (across animals, across sessions), the tracking statistics were as follows: 98.51% above the base of the tail (total range: 91-99%), 97.66% for the middle of the back (total range: 87-99%), and 96.33% for between the shoulder blades (total range: 83-99%). Each of the head markers, the mean percentages were 99% for head 1 (total range: 99-99%), 98.78% for head 2 (total range: 85-99%), 98.61% for head 3 (total range: 88-99%) and 98.89% for head 4 (total range: 95-99%). The head as a whole was captured 98.82% of the time (total range: 85-99%). After each session was labeled, remaining unlabeled markers were deleted and the data were exported as a.csv file, with units in metres and individual markers (not rigid body) as output. A brief, custom-written Python script was used to convert the.csv file into a .pkl file, which was modified for usage in a custom graphical user interface (GUI). In the GUI, the head coordinate system was constructed and the tracking data was merged with spike data for further processing.

For the remaining 4 animals (tracked using Motive version 1.8), raw data was exported as a.csv file, with units in metres and individual markers as output. The.csv file was then loaded into the GUI and each point was hand labeled for > 90% of the time it was tracked each session. The mean number of frames across all open field sessions was 160159.1 (total range: 73231-296021).

For all animals, files containing tracked points were loaded into the GUI, where the coordinate system for the rigid body on the head was translated to minimize jitter, which roughly corresponded to placing it at the base of the head. We then rotated the coordinate system such that, on average, the x-direction of the head closely matched the horizontal movement direction.

### Extracting behavioral variables from tracking data

Following the recording, we labeled tracked points within the Motive (OptiTrack) interface, and imported the labeled data into a custom script in Fiji. Using the four tracked points on the animal’s head, we estimated the geometry of the rigid body using the average pairwise distances between markers. We then found the time point at which this geometry was closest to the average, and used that time point as a template. We then assigned an XYZ coordinate system to the template with the origin located at the centroid of the four points, and constructed coordinate systems at each time point of the experiment by finding the optimal rigid body transformation^62^ of the template to the location of the head markers. In order to find the likely axis of rotation for the head (i.e. the base of the head), we found the translation of the coordinate system that minimized the Euclidean distance between the origin at time point t-20 and t+20, where t is measured in frames from the tracking system (120 Hz). Next, the coordinate system was rotated to most closely match the Z-direction with the vertical direction of the room, and X-direction with that of the running direction, which was defined by horizontal movements of the origin from t-50 to t+50. Only time points where speed exceeded 10 cm/s were used to estimate running direction. The two objectives were combined by considering the sum of squared differences of the two sets of angles. This definition of running direction was used only to rotate the head direction, and was not used in subsequent analyses. Hyperparameters were chosen such that head placement using the resulting coordinate system visibly matched experiments.

To compute the variables for relating tracking to neural activity, we first denoted body direction as the horizontal component of the vector from the marker on the animal’s rear to the neck point. The angles of the head (pitch, azimuth and roll) relative to body direction were then computed assuming the XYZ Euler angle method. The back angles (pitch and azimuth) were determined relative to body direction using standard 2D rotations, which were optimally rotated such that the average peak of the occupancy was close to zero. The point on the neck was then used to determine neck elevation relative to the floor, as well as the horizontal position of the animal in the environment. Movement variables were estimated from the tracked angles using a central difference derivative with a time offset of 10 bins. Running speed was then estimated using a moving window of radius 15 bins. The values for self-motion were computed as the speed of the animal multiplied by the X and Y component of the difference in angles between the body direction at t-15 and t+15. We found this a more reliable representation of self-motion compared to previous mathematical descriptions^15^.

### 1D tuning curves

Angular behavioral variables were binned in 5°, with exception of back angles which were lowered to 2.5°. Movement variables were binned in 36 equally-spaced bins, spanning the range of recorded variables such that there was a minimum occupancy of 400 milliseconds in both the first and last bins. Neck elevation bins were 1cm, while position in the environment was estimated using 6.67cm bins. Finally, self-motion used a bin size of 3cm/s. For all rate maps, the average firing rate per bin was calculated as the total number of spikes per bin, divided by total time spent in the bin. All smoothed rate maps were constructed with a Gaussian filter with standard deviations of 1 bin. Only bins with a minimum occupancy of 400 milliseconds were used for subsequent analysis. To compare with shuffled distributions, we shifted the neural activity 1000 times on the interval of ±[15,60] seconds.

### Model selection

To determine which of the large number of tracked features best explained the neural activity we considered the neural activity binned to match the resolution of the tracking system (120 Hz). We fit the neural data using a Bernoulli generalized linear model^63^ assuming different versions of the natural parameter for each neurons, where each version contained a feature or sets of features as well as the constant term. We included each of the six postural features, their derivative values, body direction, head direction, speed, position and self-motion each in single variable models. We then maximized the likelihood of the data given each of the models using an L1 regularizer with λ= 10^−4^ for each model across 10 folds of the data. We tested all combinations of angles associated with the head, back, and a model with all six behavioral variables.

As in Hardcastle et al. (2017), we first compared the single feature models using the average cross-validated log-likelihood ratio, keeping the model, if any, that had the highest positive score and was significantly different from zero. Significance was determined using a one-sided signed rank test with significance of 0.01. Additional features were included if the increase in cross-validated log-likelihood ratio of the more complicated model was significantly more than that of the less-complicated model. Significance was again determined using a one-sided signed rank test with significance value of 0.01.

### Decoding

In order to visualize and decode in the space of the six posture variables we used a common dimensionality reduction technique, Isomap^36^, to reduce the space to two dimensions. We did this by first normalizing each of the features, dividing the values for each by the maximum of the absolute value of the feature. This bounded the angular values to [-1,1] and the neck elevation to [0,1]. We then applied Isomap from the python library Scikit-learn^64^ with an assumed number of neighbors equal to 100, which resulted in an estimated reconstruction error of 0.03.

In order to decode on this space, we again assumed the neural data were distributed according to a Bernoulli distribution and fit it with a GLM. In this case, however, we used a square grid, 22 × 22, of Gaussian functions with width of 0.25 units to serve as a basis set^65^ for the Isomap surface, spanning the total space of 2.99 × 2.53 units. We used a small L1 regularizer with λ= 10^−7^ and a 10 fold cross-validation scheme. The number and width of the basis functions as well as the value for the regularizer were determined by optimizing the average log-likelihood of the test sets. After fitting the model to the data, we computed the log-likelihood at each time point of the test data and for each bin of the binned posture map (50 × 50). To then visualize this as a dynamic map, Q(t,x,y), of the animal’s likely position (x,y) in this posture space at time t, we let Q(t,x,y)=exp(L_ave_(t,x,y)), where L_ave_(t,x,y) was determined from a moving 150ms Gaussian average of the log-likelihood of the test data for each posture bin (x,y). The maps of Q(t,x,y) can be visualized in both Supplementary videos 7-8 as well as in Figure 4a. Only bins occupied >150ms are displayed in the figures. The most likely position at a given time bin t was then determined by finding the x,y position corresponding to the peak value of Q(t,x,y). The error between the values for x and y and the true position were then computed using the Euclidean distance. This was done for the entire recording of 59 neurons as well as with subsets of neurons. To estimate performance with fewer neurons, we built the decoder using random subsets of 2, 6, 10, … 54 and 58 neurons. Decoding was repeated 100 times for each population size to compute a mean decoding accuracy. For each population size, chance values were computed by shuffling the spikes of each neuron independently by a number of ±[10,120] seconds. This procedure was repeated 100 times for each population size. To estimate the likely number of neurons required for perfect decoding of the posture space, we extrapolated a linear fit to the reconstruction error as a function of the log of the number of neurons. Finally, decoder error as a function of bins in the posture map was estimated by taking the average error of the moments the animal was located in a given posture bin.

### Cross-correlations

In sessions with dual simultaneous recordings, firing rates were calculated for each cell in 5ms bins and smoothed with a Gaussian kernel (S.D. = 100ms). Pairwise cross-correlations (Pearson’s *r*) were computed between the activity of any given M2 cell with each of the simultaneously-recorded PPC cells by offsetting the spiking activity of the PPC cell at intervals of 5ms from −5 to 5 seconds. To obtain values for the null-distribution, the first cell pair was cross-correlated 1000 times with a random offset each time, ranging from ±20 to 40 seconds. Pairs were considered to be significant if their peak cross-correlation was greater than 0.1 (in absolute terms) and if they exceeded chance values by 6 S.D. for 200ms, centered at the peak. The example pair cross-correlations were z-scored, so session 1 and 2 could be shown on the same scale. Group results were obtained by normalizing all significant cross-correlations series by the absolute value of their peak and then averaging them at every time point. The grey shaded area represents the 99% C.I. for the resulting mean values. Bootstrapped values for the peaks were generated by resampling from the cross-correlation sets 1000 times with 10 or 50% of the significant positive or negative series, respectively. For each of the bootstraps, selected series were normalized by the absolute value of their peak and then averaged at every time point. For each of the 1000 samples, we established the temporal offset of the peak/trough in the curve. We calculated the S.D. of the distribution of the peak/trough offsets and considered that the standard error of the population mean and obtained confidence intervals around bootstrapped means, which corresponded to the observed peaks to the third decimal.

### Histology

Silicon shanks were left in place after the final recording session. The rats received an overdose of Isofluorane and were perfused intracardially with saline and 10 % formalin or 4% paraformaldehyde. Electrodes were removed 60-120 min after perfusion, and brains were extracted and stored in DMSO. Frozen sagittal sections (30 μm) were cut in a cryostat, all sections were mounted on glass slides and stained with cresyl violet. Using a digital scanner and scanning software (Carl Zeiss AS, Oslo, Norway), electrode locations in PPC and M2 were registered with respect to local anatomical landmarks. Recoding sites were located on photomicrographs obtained using ZEN (blue edition) and imported to Adobe Illustrator. The position of the electrodes during recording was extrapolated using the read-out of the tetrode turning protocol and taking shrinkage (~20%) from histological procedures into account. Recording depths in PPC ranged from 500-1800μm, and in M2 ranged from 500-1800μm.

---

**Suppl. Movie 1**. Animated demonstration of the behavioral arena and recreated 3D rat as rendered by our graphical user interface (GUI).

**Suppl. Movie 2**. Overhead view of rat foraging in the open field in the GUI. The animal is held in place artificially to visualize movement of the body, and spiking activity of a single PPC neuron can be heard when the animal flexes its head and spine to the right.

**Suppl. Movie 3**. Same as Suppl. Movie 2, though in a different animal and viewed from the side, with a PPC neuron driven by rearing.

**Suppl. Movie 4**. Example of an M2 neuron selective for leftward roll of the head.

**Suppl. Movie 5**. An M2 neuron driven by the combined turn of the head to the right while the animal rears on its haunches. The neuron continues to spike while the animal remains still in the cell’s preferred body position.

**Suppl. Movie6**. A PPC neuron that fires maximally when the head is raised at a high angle relative to the back, tending to occur just before rearing.

**Suppl. Movie 7**. (**left**) 3D animation depicting the behavior of a rat in the open field, played at 1/3 speed. (**right**) Synchronized frame-by-frame decoding of the animal’s posture on a 2D “posture map” using simultaneously recorded ensembles in PPC and M2. The true posture of the rat is indicated by a green “X”, and color-coding depicts the posterior distribution of the animal’s posture estimated using spiking activity from PPC and M2.

**Suppl. Movie 8**. Same as Suppl. Movie 7, taken from later in the same recording session.

## Acknowledgements

We thank B. McNaughton, M. Long, Y. Dan and E. Moser for comments on the manuscript; K. Haugen, K. Jenssen, E. Kråkvik, H. Obenhaus, R. Gardner, T. Feyissa, K. Hovde, H. Kleven, M. Gianatti and H. Waade for technical and I.T. assistance; J. Jeon, J. Adams and NeuroNexus for assistance in drive design; G.M. Olsen and M. Witter for assistance with anatomical delineations; S. Eggen for veterinary oversight. This study was supported by research grants from the European Research Council (‘RAT MIRROR CELL’, Starting Grant Agreement N^o^ 335328), the Research Council of Norway (FRIPRO Young Research Talents, Grant Agreement N^o^ 239963), the Kavli Foundation, and the Centre of Excellence scheme of the Research Council of Norway (Centre for Neural Computation).

## Author contributions

J.W. and B.D. designed the experiments. B.M. and T.T. conducted the experiments. B.D. designed the analyses, B.D., B.M., S.B. and J.W. performed the analyses. J.W. and B.M. wrote the paper with assistance from T.T. and B.D.

---

**Extended Data Figure 1.**
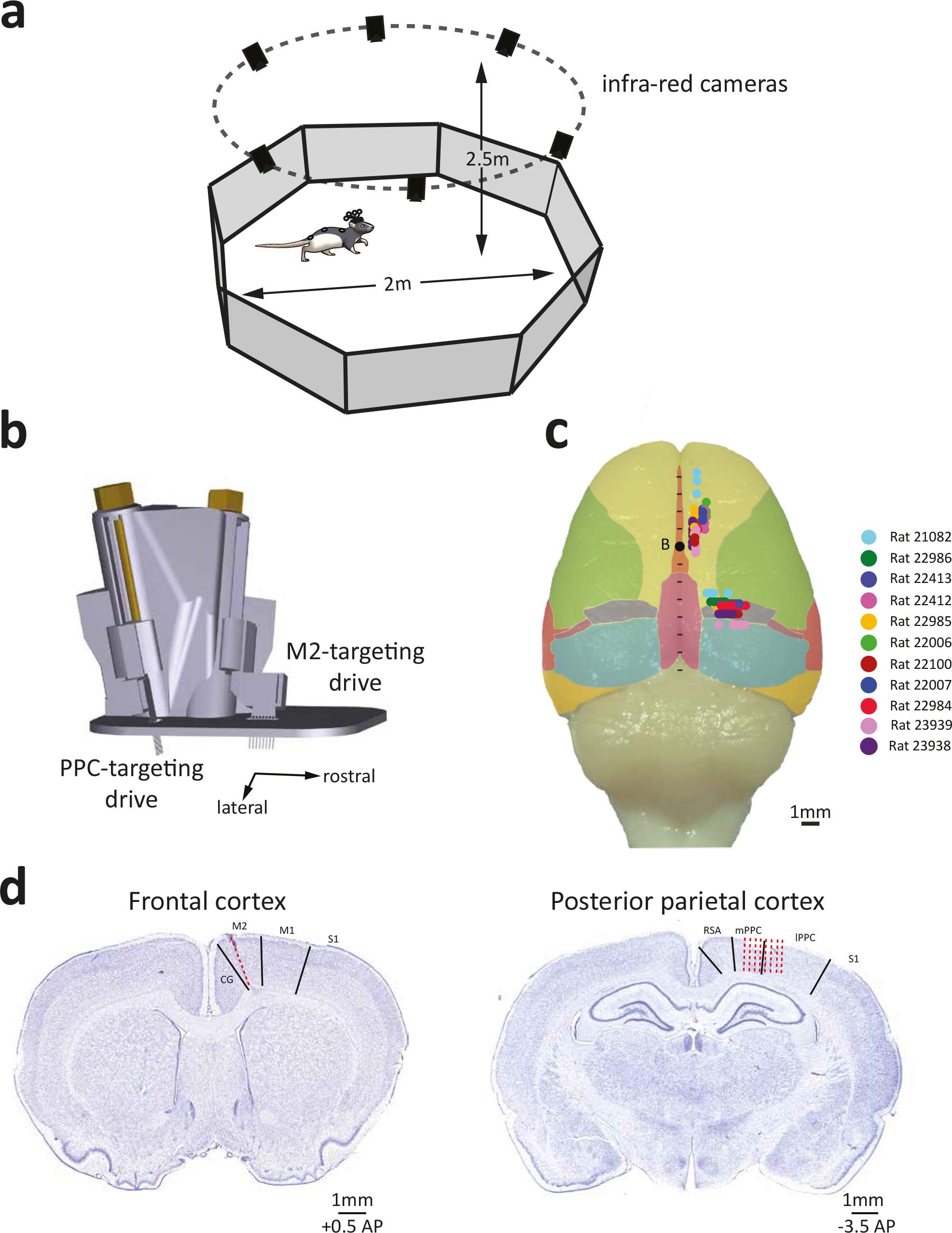
Experimental methods for 3D tracking and neural recordings. **a**, Rats marked with retrorefletive markers ran in a 2m open octagonal arena, with 6 high-speed (120fps) infrared cameras arranged above in a circle. **b**, Custom dual microdrives were used to target 8-shank silicone probes to PPC and M2. The M2 drive was designed to target the cortical surface 0.75mm lateral of midline, with tetrodes tilted 20° laterally to optimize travel in M2. The PPC drive was positioned 4mm behind the caudal-most shank on the M2 drive, and was centred 2.7mm lateral of midline. The PPC drive was tilted 10° so that shanks advanced slightly rostrally with turning. **c**, Aeral view of cortical penetrations yielding single units in 11 rats. Sites recorded with the frontal drive ranged from −0.5 AP to +4.2 AP, and from ~0.6 to 2.0mm ML. Shanks in PPC spanned 1.7 to 3.7 ML, and −3.0 to −4.5 AP, and in two animals the posterior drive penetrated into caudal M2. **d**, Cresyl violet-stained coronal sections (40μm thick) showing recording tracks (red stippled lines) in M2 (left) and PPC (right).

**Extended Data Figure 2.**
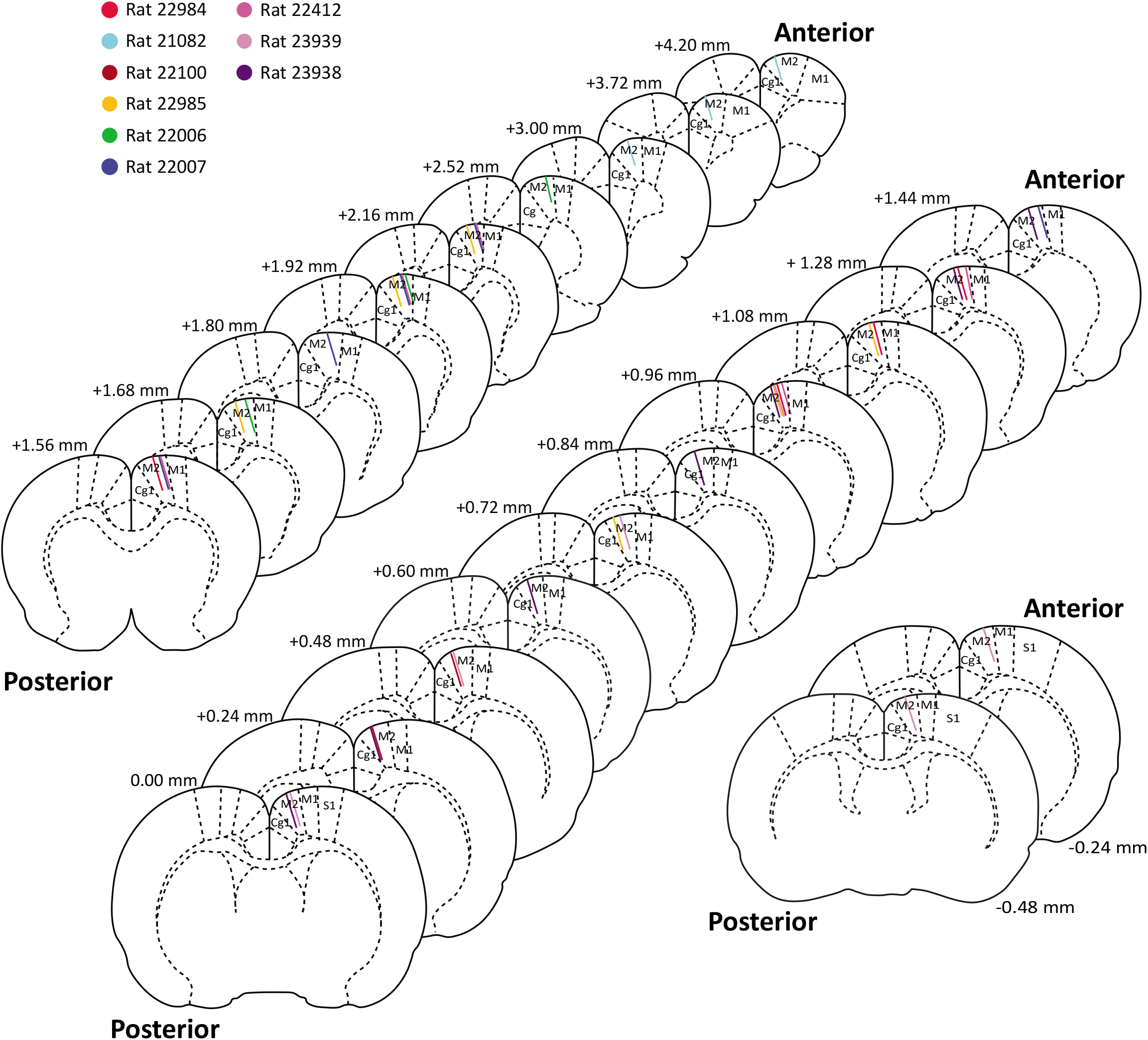
Recording locations of each shank that produced single units M2. Coronal maps showing architectonic boundaries at each AP location where single units were recorded in M2 in nine rats. Recording locations in M2 ranged from +4.20mm to −0.48mm AP; all recording sites were confirmed histologically as confined to M2.

**Extended Data Figure 3.**
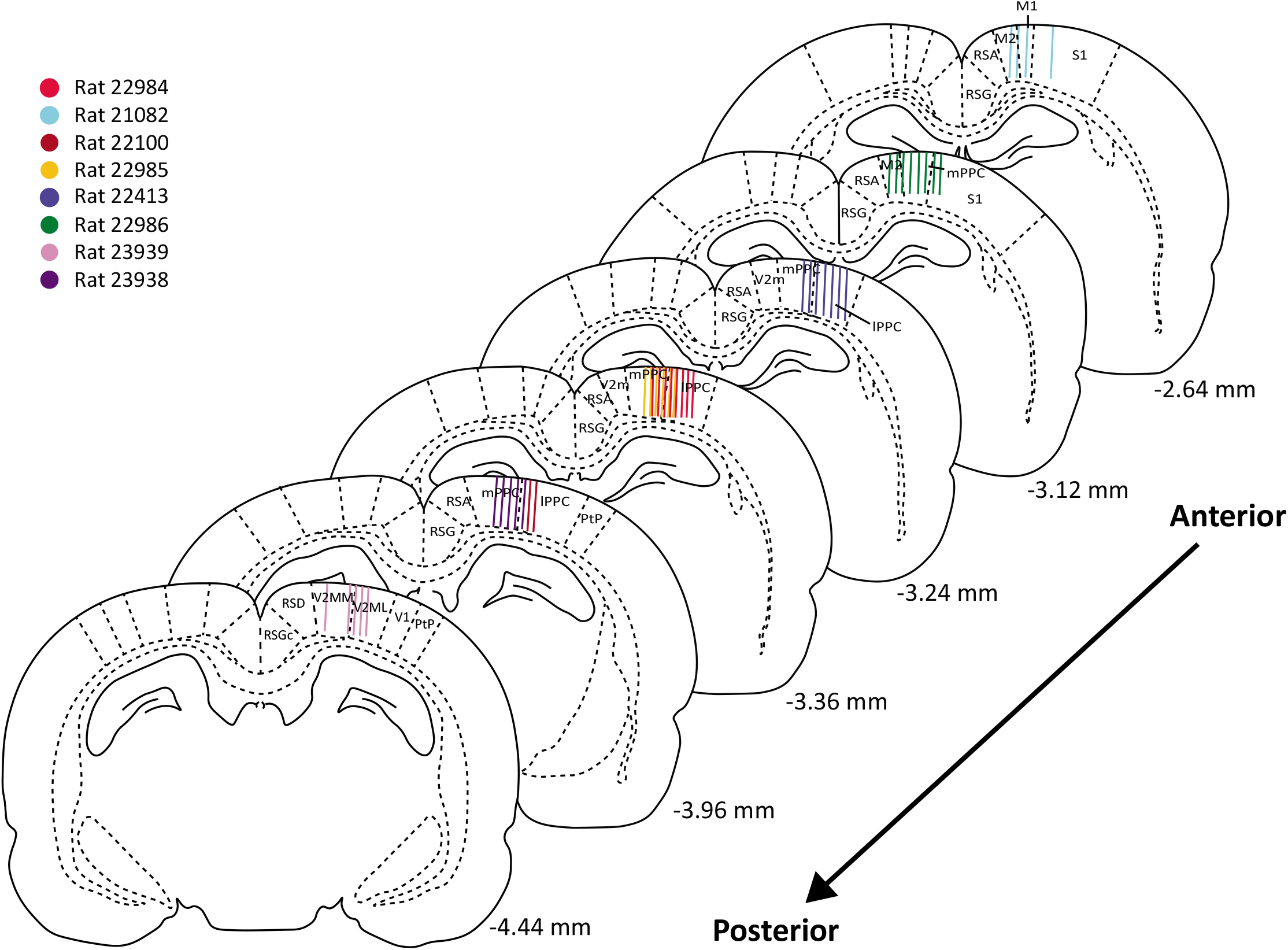
Recording locations of shanks targeting PPC. Coronal maps showing architectonic boundaries at each AP location where single units were recorded PPC and neighboring regions in eight rats. Recordings spanned the rostral-to-caudal extent of PPC, and in one animal (#23939) included area V2M (Paxinos & Watson 2006). The shanks in two rats (#21082 and #22986) were entirely rostral to PPC, with units recorded in caudal M2. Ten cells in #21082 were recorded in M1 and analysed separately.

**Extended Data Figure 4.**
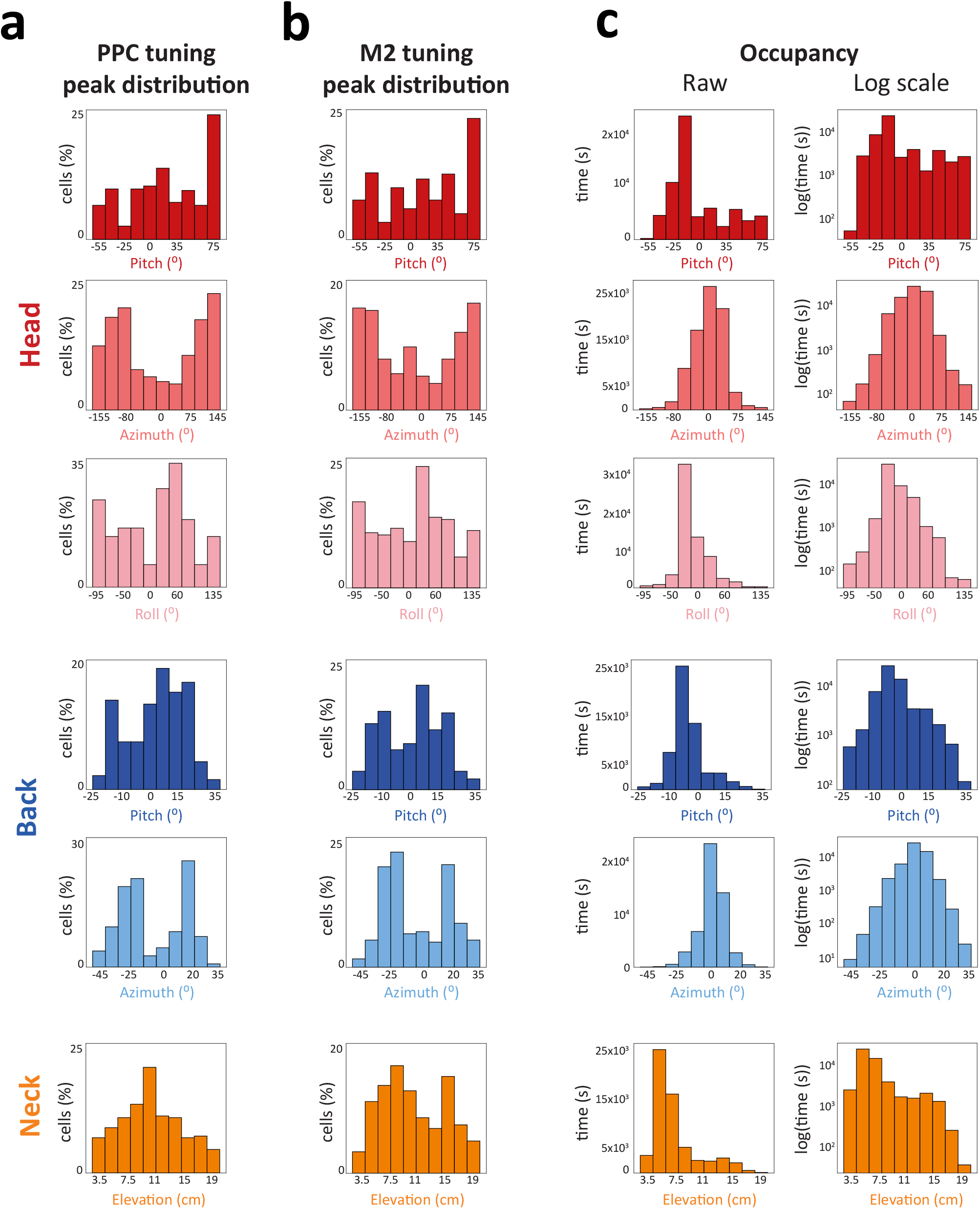
Distributions of tuning peaks in PPC and M2, and behavioural coverage. **a**, The distribution of tuning peaks for head, back and neck position in PPC. For all features tested the observed distributions of tuning peaks were significantly non-uniform (top to bottom: head pitch, χ^2^(9) = 104.3, P <0.001; head azimuth, χ^2^(9) = 64.8, P < 0.001; head roll, χ^2^(9) = 50.8, P < 0.001; back pitch, χ^2^(9) = 113.7, P < 0.001; back azimuth, χ^2^(9) = 166.6, P < 0.001; neck elevation, χ^2^(9) = 55.9, P < 0.001), tending to accumulate in bins toward the periphery and with lower cumulative occupancy. **b**, The distributions of tuning peaks in M2 were also nonuniform, tending to accrete in peripheral bins with lower occupancy (top to bottom: head pitch, χ^2^(9) = 123, P <0.001; head azimuth, χ^2^(9) = 51.5, P < 0.001; head roll, χ^2^(9) = 38.4, P < 0.001; back pitch, χ^2^(9) = 106, P < 0.001; back azimuth, χ^2^(9) = 123.5, P < 0.001; neck elevation, χ^2^(9) = 63.9, P < 0.001). c, (Left) Cumulative occupancy for all egocentric positions sampled across animals; (right) same data, but on a log scale to show that bins with the lowest occupancy had >5 sec of sampling.

**Extended Data Figure 5.**
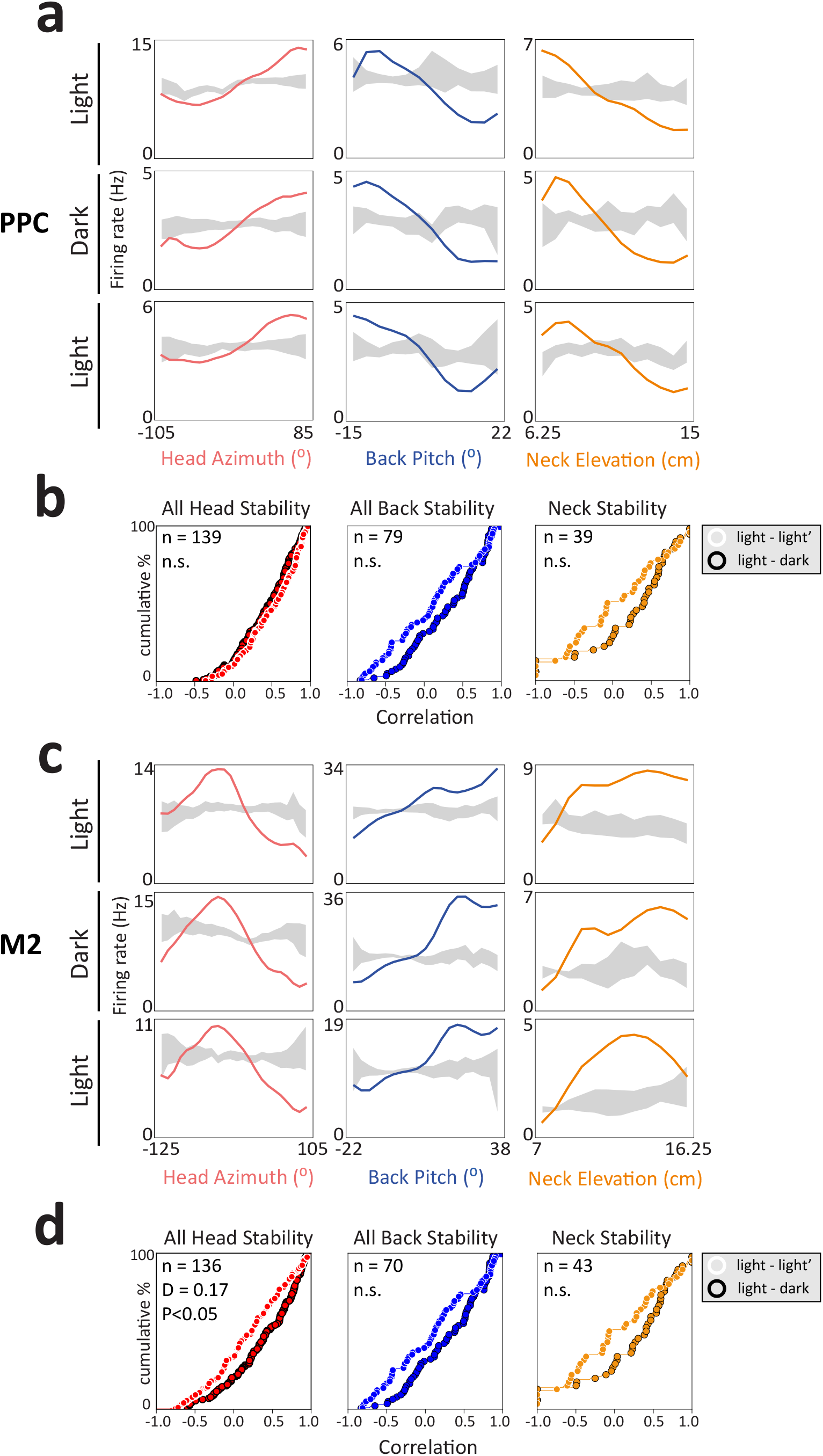
Postural tuning in PPC and M2 was stable in darkness. **a**, Examples of three cells in PPC with stable tuning to head azimuth (left), back pitch (middle) and neck elevation (right) in light and darkness. **b**, Cumulative frequency distributions of Pearson’s correlations for all features of the head (left) and back (middle), and neck elevation (right) between two sessions in light, or one light and one dark session. The distributions of correlations did not differ significantly for any comparison (inset, K-S test, p > 0.05 in all cases). **c**, Examples of three M2 cells with stable tuning across light and dark recording sessions (same as in a). **d**, Cumulative frequency distributions of Pearson’s correlations between light-light’ and light-dark recording sessions for all head (left) and back features (middle), and neck elevation (right). The stability of tuning curves for the head were the only that differed between light and dark sessions, with the light-dark session showing greater stability on average than the light-light’ session (D = 0.21, P < 0.001).

**Extended Data Figure 6.**
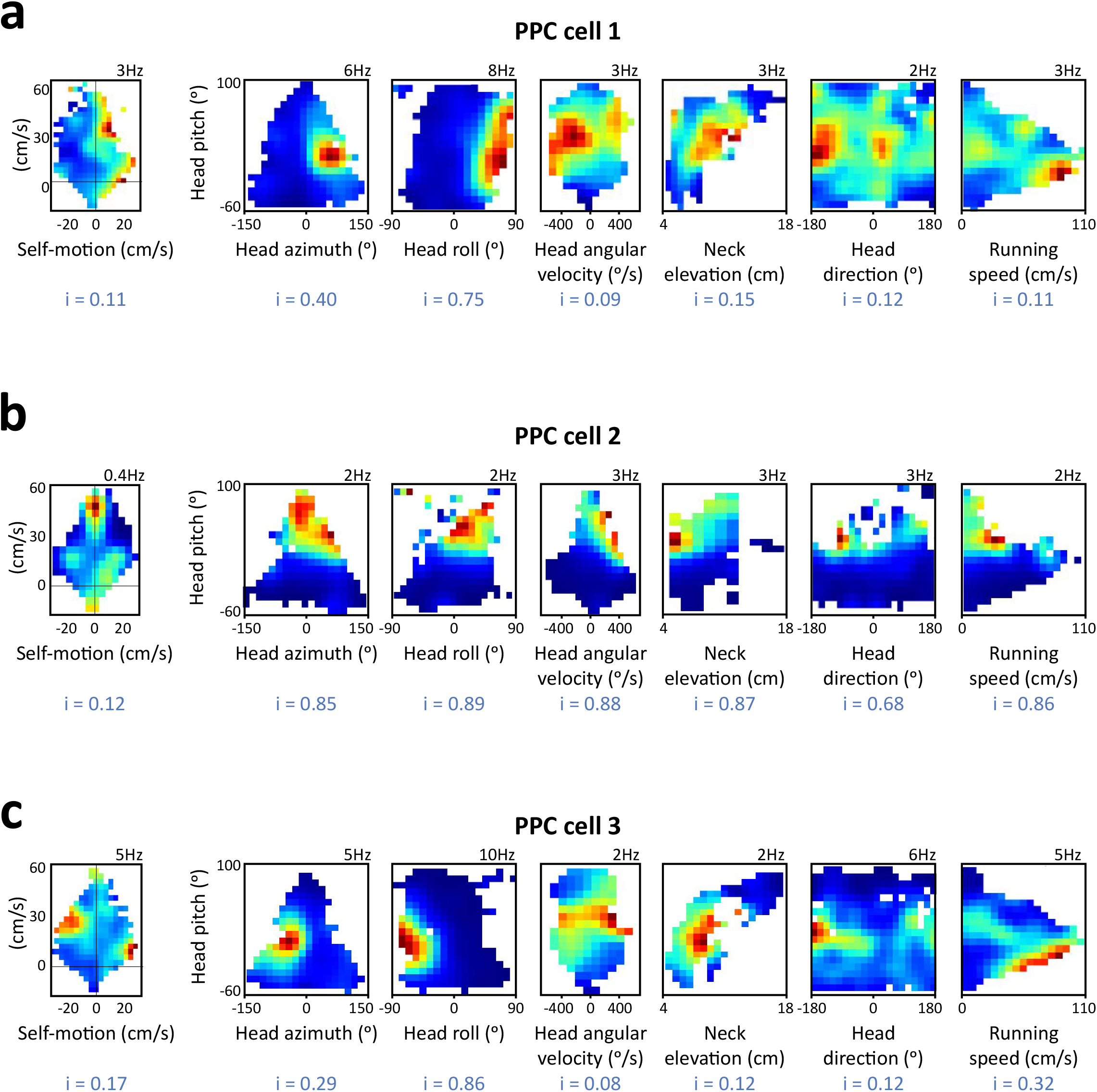
Examples of postural tuning independent of self-motion in PPC. **a**, (left) A cell showing little tuning to self-motion (information rate 0.11 bits/sec; blue text), exhibited sharp firing fields for combinations of head pitch, azimuth and roll (right, first 2 panels). The cell was largely uninformative other behavioral features, such as angular head velocity, neck elevation, head direction or running speed (reflected by lower information rates of the 2D rate maps). **b**, Example of a PPC cell with strong sensitivity to head pitch, which was not captured by the self-motion map. The cell’s selectivity for head pitch was clear in interactions with several behavioral features, including head angular velocity, neck elevation, head direction and running speed (right panels). **c**, A third cell showing little self-motion tuning that was particularly sensitive to head pitch x roll (2nd panel on right).

**Extended Data Figure 7.**
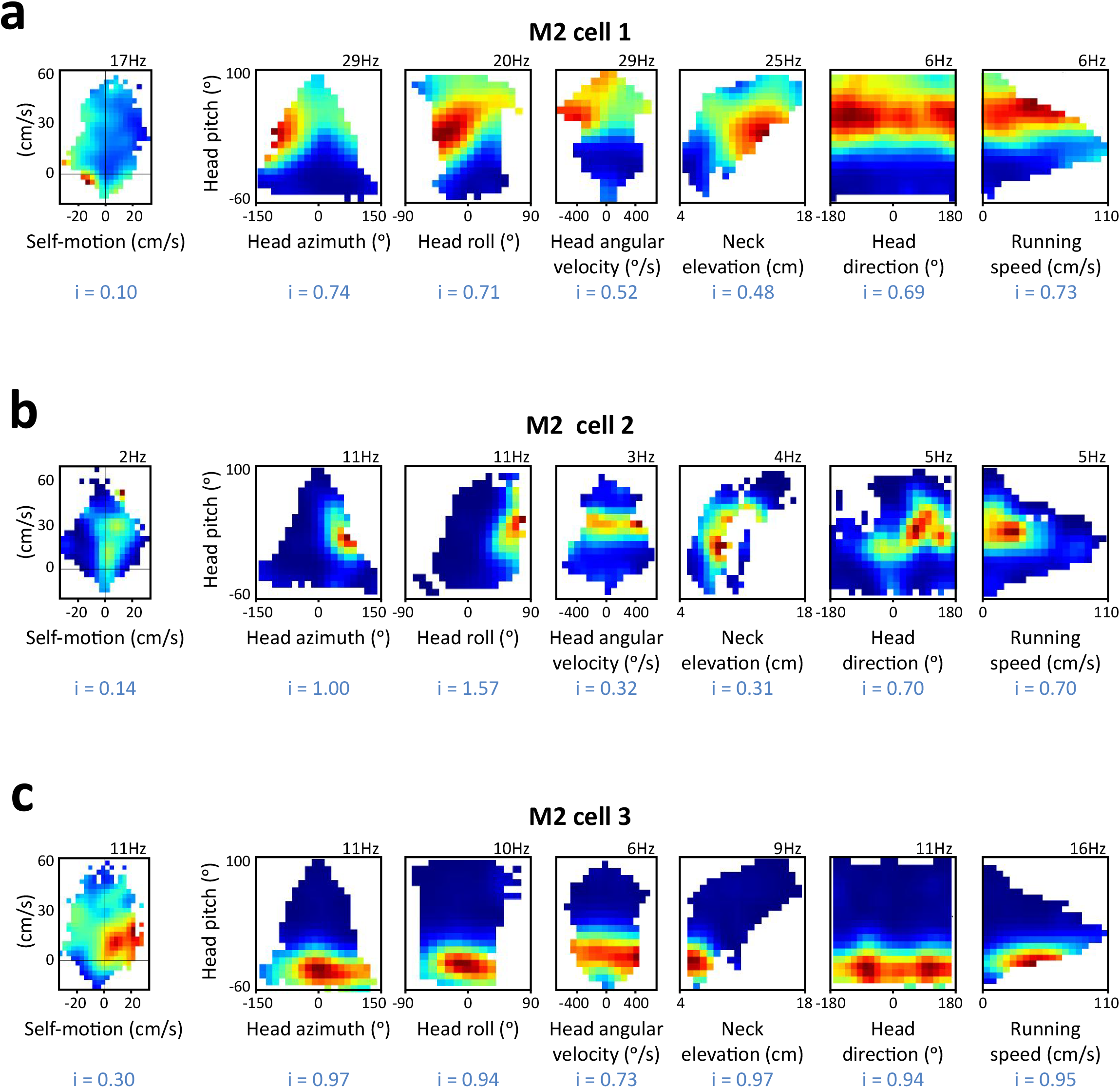
Examples postural tuning in M2 independent of self-motion. **a**, (left) Example of a cell showing poor tuning to self-motion (information rate = 0.10), but sharp sensitivity to pitch, azimuth and roll of the head (2D interaction rate maps, right). The tuning of this cell was dominated by head pitch, and showed little co-modulation by world-centred features such as head direction and speed. **b**, Another cell tuned weakly to self-motion (leftmost rate map) was highly sensitive to specific combinations of head pitch, azimuth and roll. The cell was co-modulated weakly by head direction and tended to fire at lower running speed (rightmost rate map). **c**, The firing rate of a cell showing a preference for rightward turns (self-motion map, left) is better explained by downward pitch of the head and low neck elevation. The cell did not ditinguish left from right head angular velocities middle panel), it was not sensitive to head direction, nor was it modulated by running speed. In all cases tested, 3D postural tuning was maintained when self-motion maps were split into leftward or rightward displacements (not shown).

**Extended Data Figure 8.**
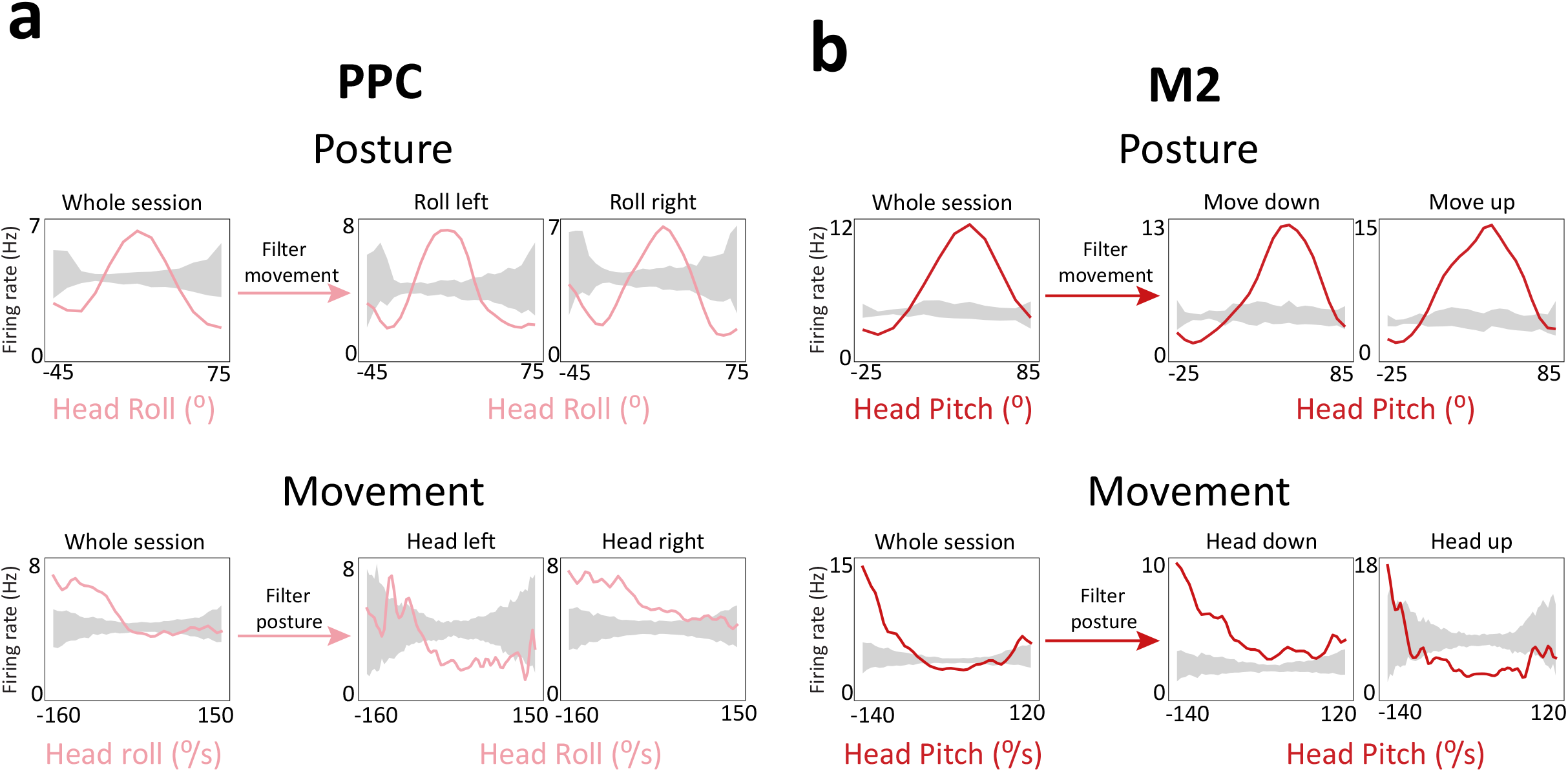
SpliШng sessions by posture or movement reveals the primacy of pose. **a**, (top, left) 1D tuning curve of a PPC neuron tuned to head roll. (top, right) The recording session was split to only include leftward or rightward angular velocities for the roll of the head, but this had no effect on the tuning curve, demonstrating that posture was independent of head movement. (below, left) The ĥ’ring rate of the same cell is expressed as a function of roll velocity. (below, right) The session was again split, but on the basis of the roll (position) of the head to the left or right. The tuning curve of the cell was not stable across these conditions (right), demonstrating that velocity tuning was downstream of postural tuning. **b**, The same test was performed on an M2 neuron tuned to head pitch, which again demonstrated that postural tuning was independent of movement velocity (top), but not *vice versa* (below).

**Extended Data Figure 9.**
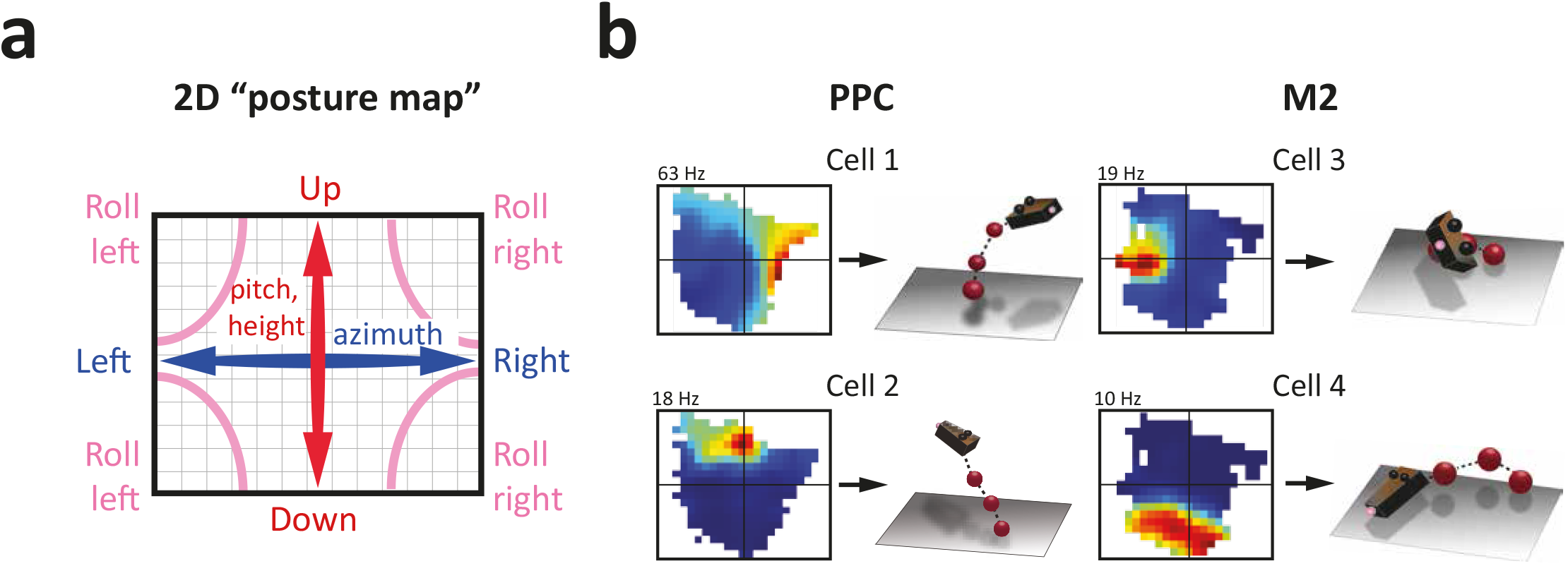
2D posture maps. **a**, The six postural features we measured (head pitch, azimuth and roll, neck elevation, back pitch and azimuth) were collapsed onto a single 2D surface using Isomap (Tenenbaum, 2000). Vertical features, such as pitch of the head, back, and neck height are stretched along the ordinate (red arrow); head and back azimuth are represented along the abscissa (blue arrow); head roll occupies the corners. **b**, Two example PPC cells (left) and two M2 cells (right) show discrete firing fields on 2D posture maps with illustrations of their corresponding 3D poses. Population-level activity was used to decode dynamic posture using Isomaps in Figure 4a.

**Extended Data Figure 10.**
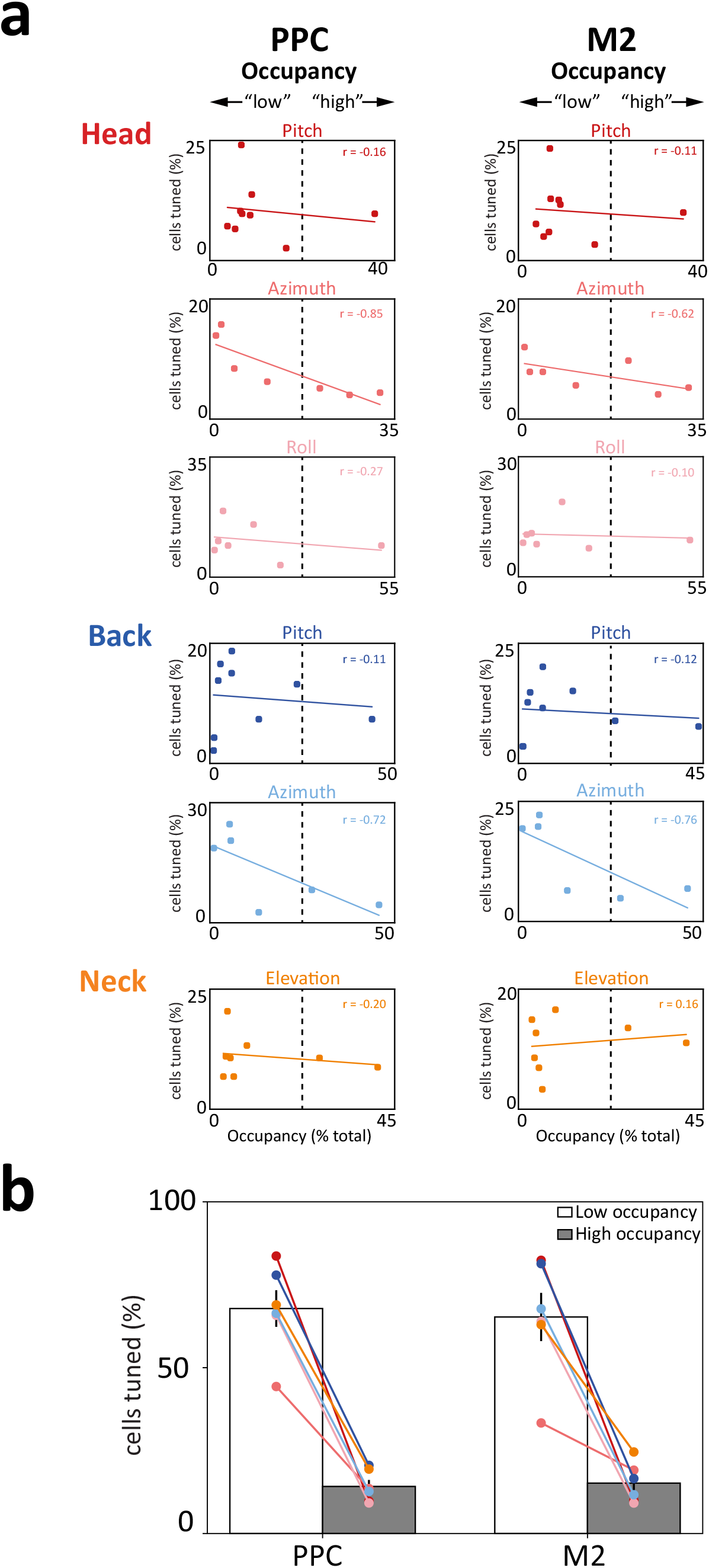
For all features, the majority of cells had tuning peaks at positions with <50% occupancy.**a** The duration that each bin was occupied for measures of head, neck and back position (as shown in Extended Data Fig.4 a and b) was divided by total recording time to determine the percent occupancy per bin. Bins with <1% occupancy were excluded. The percentage of cells with significant tuning peaks (>3 S.D. above shuffled) was higher for posture bins with below-mean occupancy (e.g. for head pitch in both PPC and M2, 83% of cells had tuning peaks in bins with less than 20% relative occupancy). The inverse relationship between the number of tuned cells and occupancy was particularly clear for head and back azimuth. b, Summary of results obtained in (a) with means and SEMs. The percentages of cells tuned were summed across levels of high/low occupancy in both regions, showing that that the mean % of cells tuned for “low occupancy” in PPC was 68% (99% C.I.: 53-82%), while the mean % for “high occupancy” was 14% (99% C.I.: 9-19%). For M2, the mean % for low occupancy was 65% (99% C.I.: 47-84%), and 15% (99% C.I.: 9-22%) for “high occupancy.”

**Extended Data Table 1.** Stability of 1D tuning curves

| PPC |  |  |  | M2 |  |  |
| --- | --- | --- | --- | --- | --- | --- |
| Feature | # cells<sup>1</sup> | # stable cells<sup>2</sup> | % stable<sup>3</sup> | # cells | # stable cells | % stable |
| Head pitch | 296 | 191 | 64.5% | 341 | 218 | 63.9% |
| Head azimuth | 233 | 128 | 54.9% | 208 | 117 | 56.2% |
| Head roll | 191 | 137 | 71.7% | 236 | 156 | 66.1% |
| Back pitch | 277 | 121 | 43.6% | 269 | 140 | 52.0% |
| Back azimuth | 220 | 113 | 51.3% | 190 | 87 | 45.7% |
| Neck elevation | 260 | 143 | 55.0% | 286 | 167 | 58.4% |

<sup>3</sup>(Column2 / column 1) x 100.

^1^Number of cells with tuning curve peaks exceeding the 99^th^ percentile of the shuffled distribution in the first recording session. All cells from PPC (n = 729) and M2 (n = 808) were considered for each feature. ^2^Number of cells from the first column with across-session r-value exceeding the 95^th^ percentile of the shuffled data. ^3^(Column2 / colum 1) × 100.

**Extended Data Table 2.**
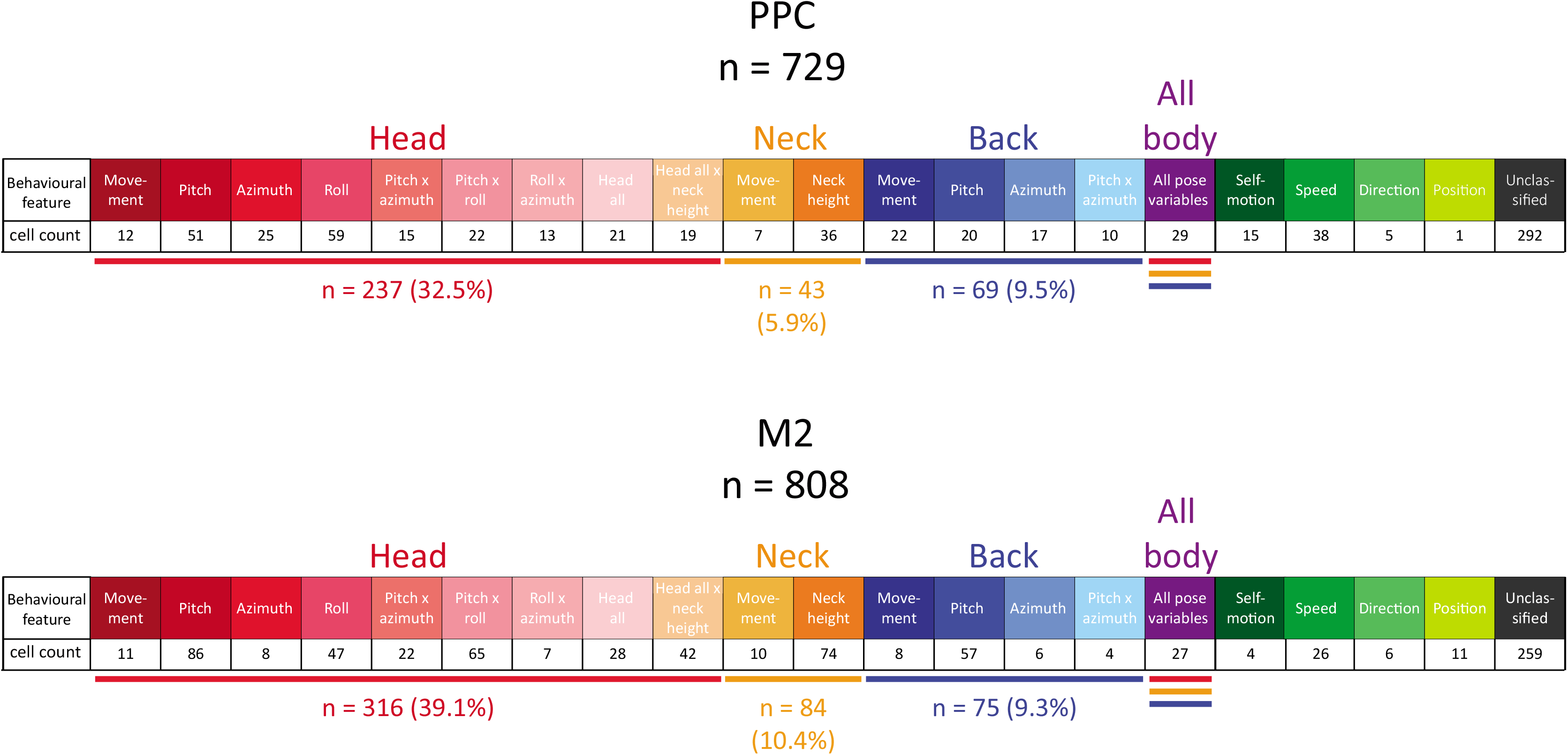
Summary of GLM model selection for all cells

